# Eukaryotic antiviral immune proteins arose via convergence, horizontal transfer, and ancient inheritance

**DOI:** 10.1101/2023.06.27.546753

**Authors:** Edward M. Culbertson, Tera C. Levin

**Affiliations:** University of Pittsburgh, Department of Biological Sciences

**Author notes:** Address correspondence to Tera C. Levin.

## Abstract

Animals use a variety of cell-autonomous innate immune proteins to detect viral infections and prevent replication. Recent studies have discovered that a subset of mammalian antiviral proteins have homology to anti-phage defense proteins in bacteria, implying that there are aspects of innate immunity that are shared across the Tree of Life. While the majority of these studies have focused on characterizing the diversity and biochemical functions of the bacterial proteins, the evolutionary relationships between animal and bacterial proteins are less clear. This ambiguity is partly due to the long evolutionary distances separating animal and bacterial proteins, which obscures their relationships. Here, we tackle this problem for three innate immune families (CD-NTases [including cGAS], STINGs, and Viperins) by deeply sampling protein diversity across eukaryotes. We find that Viperins and OAS family CD-NTases are truly ancient immune proteins, likely inherited since the last eukaryotic common ancestor and possibly longer. In contrast, we find other immune proteins that arose via at least four independent events of horizontal gene transfer (HGT) from bacteria. Two of these events allowed algae to acquire new bacterial viperins, while two more HGT events gave rise to distinct superfamilies of eukaryotic CD-NTases: the Mab21 superfamily (containing cGAS) which has diversified via a series of animal-specific duplications, and a previously undefined eSMODS superfamily, which more closely resembles bacterial CD-NTases. Finally, we found that cGAS and STING proteins have substantially different histories, with STINGs arising via convergent domain shuffling in bacteria and eukaryotes. Overall, our findings paint a picture of eukaryotic innate immunity as highly dynamic, where eukaryotes build upon their ancient antiviral repertoires through the reuse of protein domains and by repeatedly sampling a rich reservoir of bacterial anti-phage genes.

## Introduction

As the first line of defense against pathogens, all forms of life rely on cell-autonomous innate immunity to recognize threats and respond with countermeasures. Until recently, many components of innate immunity were thought to be lineage-specific[1]. However, new studies have revealed that an ever-growing number of proteins used in mammalian antiviral immunity are homologous to bacterial immune proteins used to fight off bacteriophage infections. This list includes Argonaute, CARD domains, cGAS and other CD-NTases, Death-like domains, Gasdermin, NACHT domains, STING, SamHD1, TRADD-N domains, TIR domains, and Viperin, among others[2–13]. This discovery has been surprising and exciting, as it implies that some cellular defenses have deep commonalities spanning across the entire Tree of Life. But despite significant homology, these bacterial and animal immune proteins are often distinct in their molecular functions and operate within dramatically different signaling pathways (reviewed here[5]). *How, then, have animals and other eukaryotes acquired these immune proteins?*

One common hypothesis in the field is that these immune proteins are ancient, and have been inherited since the last common ancestor of bacteria and eukaryotes[5]. In other cases, horizontal gene transfer (HGT) between bacteria and eukaryotes has been invoked to explain the similarities[6,14]. However, because most papers in this field have focused on searching genomic databases for new bacterial immune genes and biochemically characterizing them, the evolution of these proteins in eukaryotes has not been as thoroughly investigated.

To address this knowledge gap, we turned to the EukProt database, which has been specifically developed to reflect the true scope of eukaryotic diversity through the genomes and transcriptomes of nearly 1,000 species, specifically selected to span the eukaryotic tree [15]. EukProt contains sequences from NCBI and Ensemble, plus many diverged eukaryotic species not found in any other database, making it a unique resource for eukaryotic diversity[15]. While it can be challenging to acquire diverse eukaryotic sequences from traditional databases due to an overrepresentation of metazoan data[16], EukProt ameliorates this bias by downsampling traditionally overrepresented taxa.

Using this database, we investigated the ancestry of three gene families that are shared between animal and bacterial immunity: Stimulator of Interferon Gamma (STING), cyclic GMP-AMP synthase (cGAS) and its broader family of cGAS-DncV-like nucleotidyltransferases (CD-NTases), and Viperin. STING, CD-NTases, and Viperin are all interferon-stimulated genes that function as antiviral immune modules, disrupting the viral life cycle by activating downstream immune genes, sensing viral infection, or disrupting viral processes, respectively[17]. We found eukaryotic CD-NTases arose following multiple HGT events between bacteria and eukaryotes. cGAS falls within a unique, mainly metazoan clade. In contrast, OAS-like proteins were independently acquired and are the predominant type of CD-NTase found across most eukaryotes. Separately, we have discovered diverged eukaryotic STING proteins that bridge the evolutionary gap between metazoan and bacterial STINGs, as well as two separate instances where bacteria and eukaryotes have acquired similar proteins via convergent domain shuffling. Finally, we find that Viperin is likely to be truly ancient, with both broad representation across the eukaryotic tree of life and evidence of two additional HGT events where eukaryotes recently acquired new bacterial viperins. Overall, our results demonstrate that immune proteins shared between bacteria and eukaryotes are evolutionarily dynamic, with eukaryotes taking multiple routes to acquire and deploy these ancient immune modules.

## Results

### Discovering immune homologs across the eukaryotic tree of life

The first step to understanding the evolution of CD-NTases, STINGs, and viperins was to acquire sequences for these proteins from across the eukaryotic tree. To search for diverse immune homologs, we employed a hidden Markov model (HMM) strategy, which has high sensitivity, a low number of false positives, and the ability to separately analyze multiple (potentially independently evolving) domains in the same protein[18–20]. To broaden our searches from initial animal homologs to eukaryotic sequences more generally, we used iterative HMM searches of the EukProt database, incorporating the hits from each search into the subsequent HMM. After using this approach to create pan-eukaryotic HMMs for each protein family, we then added in bacterial homologs to generate universal HMMs (Fig. 1A and Supp. Fig. 1), continuing our iterative searches until we either failed to find any new protein sequences or began finding proteins outside of the family of interest (Supp. Fig. 1).

**Figure 1:**
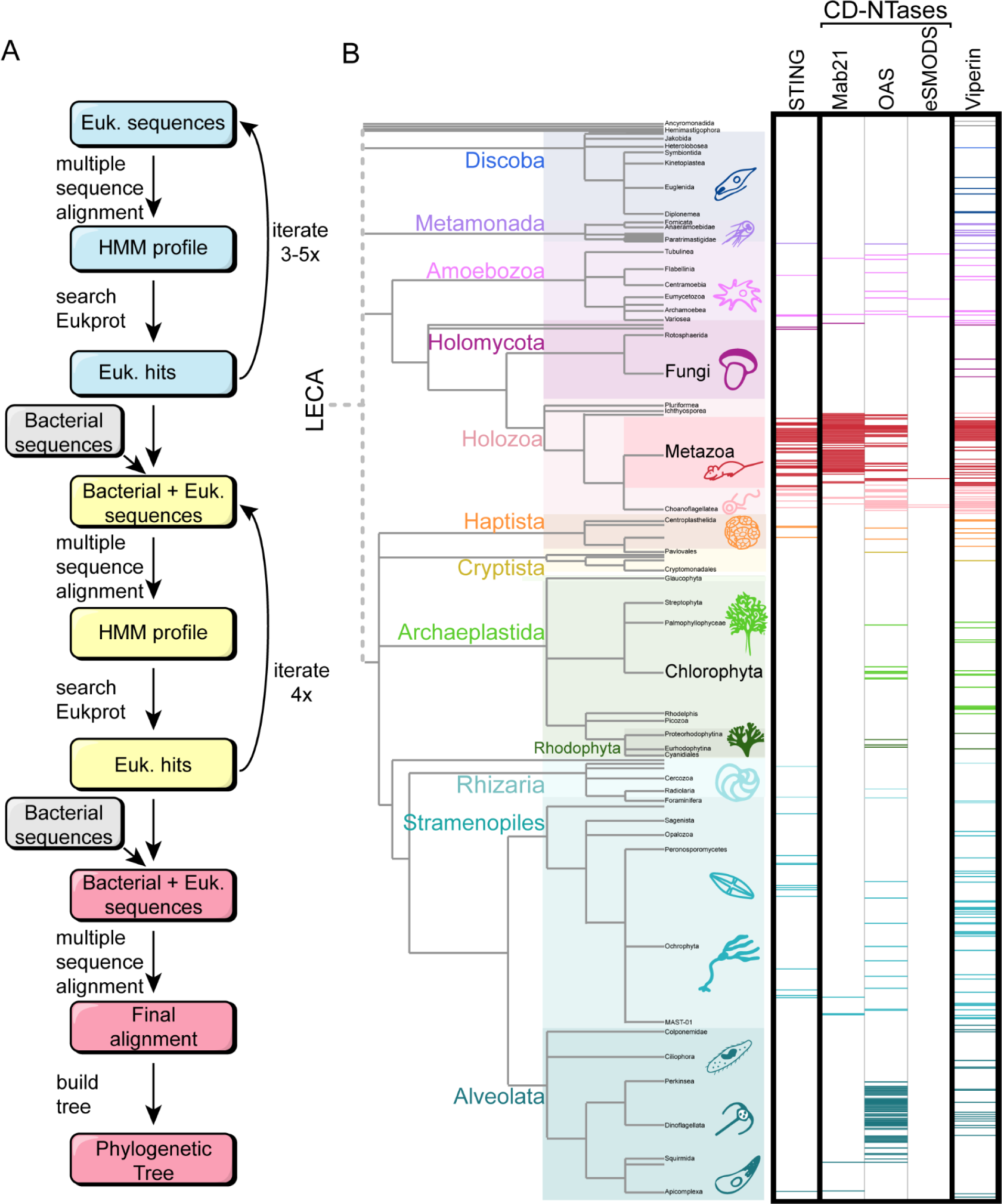
HMM searches to find homologs across the eukaryotic Tree of Life. (A) A schematic of the HMM search process. Starting from initial, animal-dominated HMM profiles for each protein family, we used iterative HMM searches of the EukProt database to generate pan-eukaryotic HMMs. These were combined with bacterial sequences to enable discovery of bacteria-like homologs in eukaryotes. Each set of searches was repeated 3-5 times until few or no additional eukaryotic sequences were recovered. (B) Phylogenetic tree of eukaryotes, with major supergroups color-coded. The height of the colored rectangles for each group is proportional to its species representation in EukProt. Horizontal, colored bars mark each eukaryotic species in which we found homologs of STINGs, CD-NTases, or Viperins. White space indicates species where we did not recover any homologs. The CD-NTase hits are divided into the three eukaryotic superfamilies defined in Fig. 2.

Our searches for CD-NTases, STINGs, and viperins recovered hundreds of eukaryotic proteins from each family, including a particularly large number of metazoan sequences (red bars, Fig. 1B). It is not surprising that we found so many metazoan homologs, as each of these proteins was discovered and characterized in metazoans and these animal genomes tend to be of higher quality than other taxa (Supp. Fig 2). We also recovered homologs from other species spread across the Eukaryotic tree, demonstrating that our approach could successfully identify deeply diverged homologs (Fig. 1B). However, outside of Metazoa, these homologs were sparsely distributed, such that for most species in our dataset (711/993), we did not recover proteins from the three immune families examined (white space, lack of colored bars, Fig. 1B). We believe this pattern reflects a pattern of ongoing, repeated gene losses across eukaryotes, as has been found for other innate immune proteins[21–23] and other types of gene families surveyed across eukaryotes[22,24–26]. We found that BUSCO completeness scores and data type (genomes vs. transcriptomes) were insufficient to explain the pattern of gene loss (Supp. Fig. 2). Thus, although it is always possible that our approach has missed some homologs, we believe the resulting data represents a fair assessment of the true diversity across eukaryotes. To analyze these genes, we aligned the homologs with MAFFT and MUSCLE and then generated phylogenetic trees with IQtree, FastTree, and RaxML-ng (see Materials and Methods). We considered our results to be robust if they were concordant across the majority of six trees generated per gene.

### Eukaryotes acquired CD-NTases from bacteria through at least three independent HGT events

We next studied the evolution of the innate immune proteins, beginning with cGAS and its broader family of CD-NTase enzymes, which generate diverse oligonucleotides. In addition to the well-studied cGAS, a number of other eukaryotic CD-NTases have been previously described: 2’-5’-Oligoadenylate Synthetase 1/2/3 (OAS1/2/3), Male abnormal 21-Like 1/2/3/4 (MAB21L1/2/3/4), Mab-21 domain containing protein 2 (MB21D2), Mitochondrial dynamics protein 49/51(MID49/51), and Inositol 1,4,5 triphosphate receptor-interacting protein 1/2 (ITPRILP/1/2)[27]. Of these, cGAS and OAS1 are the best characterized and both play roles in immune signaling. cGAS, Mab21L1, and MB21D2 are all cGAS-like receptors (cGLRs), and recent work has shown that cGLRs are present in nearly all metazoan taxa and generate diverse cyclic dinucleotide signals[28]. However, the immune functions of Mab21L1 and MB21D2 remain unclear, although they have been shown to be important for development[29–31].

Following infections or cellular damage, cGAS binds cytosolic DNA and generates cyclic GMP-AMP (cGAMP)[32–35], which then activates downstream immune responses via STING [34,36–38]. OAS1 synthesizes 2’,5’-oligoadenylates which bind and activate Ribonuclease L (RNase L)[39]. Activated RNase L is a potent endoribonuclease that degrades both host and viral RNA species, reducing viral replication (reviewed here[40,41]). Some bacterial CD-NTases such as *DncV* behave similar to animal cGAS; they are activated by phage infection and produce cGAMP[8,42,43]. Other bacterial CD-NTases generate a wide variety of dinucleotides, cyclic trinucleotides, and cyclic oligonucleotides[11]. These CD-NTases are commonly found within cyclic oligonucleotide-based anti-phage signaling systems (CBASS) across many bacterial phyla and even archaea[8,27,43].

To understand the evolutionary history of CD-NTases we used the Pfam domain PF03281 as a eukaryotic starting point. As representative bacterial CD-NTases, we used 6,132 bacterial sequences, representing a wide swath of CD-NTase diversity[43]. Following our iterative HMM searches, we recovered 313 sequences from 109 eukaryotes, of which 34 were metazoans (Supplemental Data and Fig. 1B). Most eukaryotic sequences clustered into one of two distinct superfamilies, which we name here for their highest scoring PFAM domain: Mab21 (Mab21: PF03281) or OAS (OAS1-C: PF10421) (Fig. 2A). Bacterial CD-NTases typically had sequences matching the HMM for the Second Messenger Oligonucleotide or Dinucleotide Synthetase domain (SMODS: PF18144).

**Figure 2:**
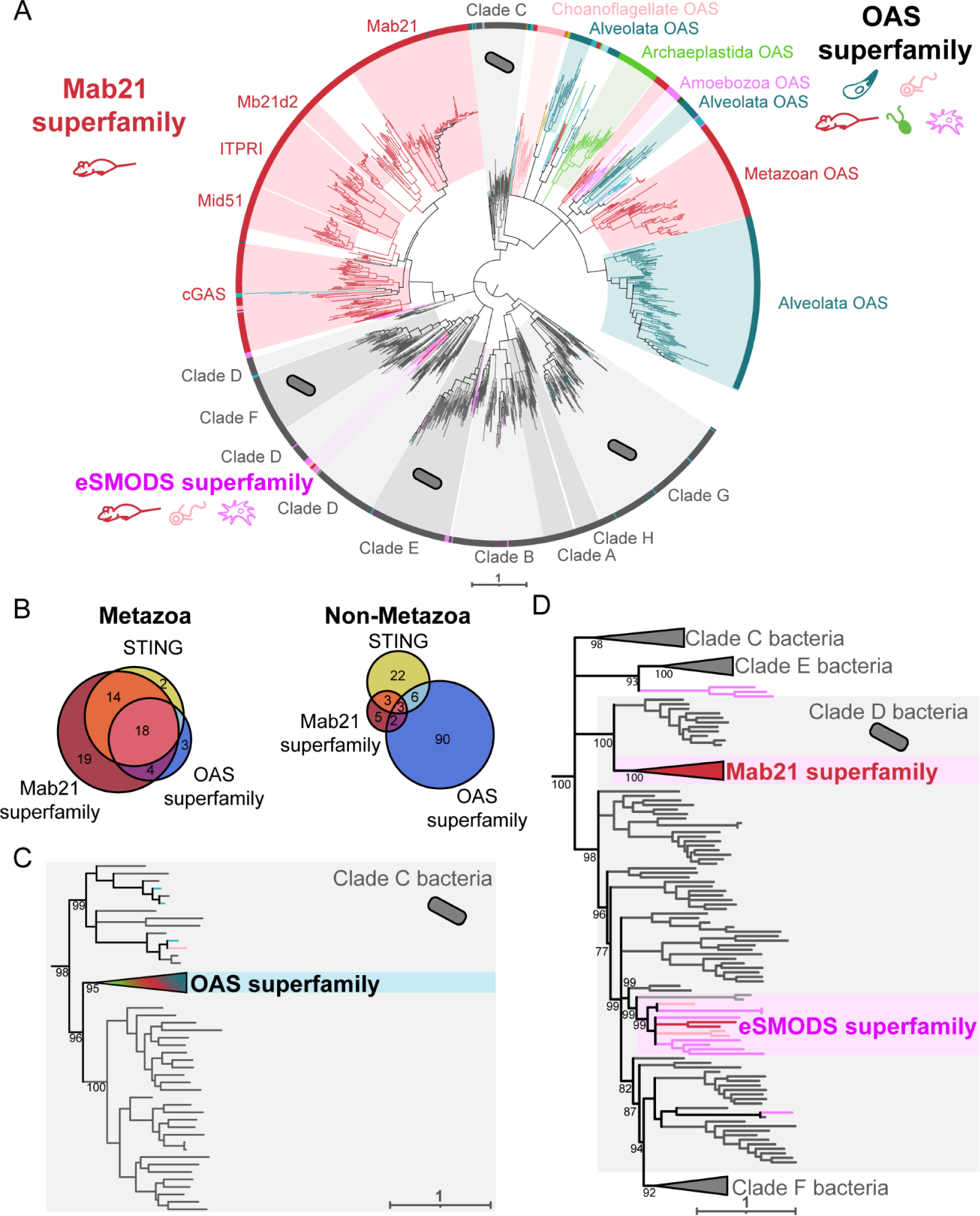
Independent HGT events gave rise to multiple CD-NTase superfamilies. (A) Maximum likelihood phylogenetic tree generated by IQtree of CD-NTases spanning eukaryotic and bacterial diversity. The Mab21 superfamily (red, top left) is largely an animal-specific innovation, with many paralogs including cGAS. In contrast, most other eukaryotic lineages encode CD-NTases from the OAS superfamily (multicolor, top right). The relatively small eSMODS superfamily (pink, bottom left) is a recent HGT between clade D bacteria and eukaryotes. Bacterial CD-NTase sequences shown in gray. Eukaryotic sequences are colored according to eukaryotic group as in Fig. 1. The tree is arbitrarily rooted on a branch separating clades A, B, G, and H, which did not typically have associated eukaryotic sequences, from the rest of the bacterial CD-NTases. (B) Venn diagrams showing the number of species where we detected at least one homolog of STING, Mab21 superfamily CD-NTases, and/or OAS superfamily CD-NTases in Metazoa (left) or non-metazoan eukaryotes (right). (C) Within the CD-NTase phylogenetic tree in A, the OAS superfamily branches within clade C bacterial CD- NTases (gray branches). (D) Clade D CD-NTases (gray branches) have been horizontally transferred into eukaryotes multiple times, giving rise to both the Mab21 superfamily and the eSMODS superfamily. Ultrafast bootstraps determined by IQtree shown at key nodes. See Supplementary Figure 4 for full CD-NTase phylogenetic tree.

The Mab21 superfamily is composed almost entirely of metazoan sequences, with only a few homologs from Amoebozoa, choanoflagellates, and other eukaryotes (Fig. 2A). Indeed, the majority of animal CD-NTases (cGAS, Mid51, Mab21, Mab21L1/2/3/4, Mb21d2, ITPRI) are paralogs within the Mab21 superfamily, which arose from repeated animal-specific duplications[44] (Supp. Fig 6). In contrast, unlike the animal-dominated Mab21 superfamily, the OAS superfamily spans a broad group of eukaryotic taxa, with OAS-like homologs present in 8/12 eukaryotic supergroups. This distribution makes OAS proteins the most common CD-NTases found across eukaryotes and implies that they arose very early in eukaryotic history, possibly within the last eukaryotic common ancestor (LECA).

Given the connections between cGAS and STING in both animals and some bacteria[3,43,45], we asked whether species that encode STING also have Mab21 and/or OAS proteins. Because the Mab21 superfamily is largely animal-specific, we performed this analysis separately in either Metazoa or with all non-metazoan eukaryotes (Fig. 2B). In animal species where we found a STING homolog, we also typically found Mab21 superfamily sequence (32/34), and a cGAS homolog in (26/34) species (Fig. 2B), consistent with the consensus that these proteins are functionally linked. We also observed 19 metazoan species that had a Mab21-like sequence with no detectable STING homolog. Almost half of these species (10/19) were arthropods, agreeing with prior findings of STING sparseness among arthropods[45]. Outside of animals, we found that species with a STING homolog typically did not have a detectable CD-NTase protein from either superfamily (22/34). While it remains possible that these STING proteins function together with a to-be-discovered CD-NTase that was absent from our dataset, we therefore hypothesize that many eukaryotes outside of metazoans and their close relatives[46] use STING and CD-NTase homologs independently of each other.

What was the evolutionary origin of eukaryotic CD-NTases? Interestingly, the Mab21 and OAS superfamilies are only distantly related to one another. Each lies nested within a different, previously defined, bacterial CD-NTase clade (Fig. 2 C and D). The OAS superfamily falls within bacterial Clade C (with the closest related bacterial CD-NTases being those of subclade C02-C03, Fig. 2C), while the metazoan Mab21 superfamily lies within bacterial Clade D (subclade D12) (Fig. 2D). We note that in this tree (Fig. 2D), Clade D does not form a single coherent clade, as was also true in the phylogeny that originally defined the bacterial CD-NTase clades [11].

We also observed a number of eukaryotic sequences scattered across different bacterial CD-NTase clades (Fig. 2A, colored branches within gray clades). While some of these may reflect additional HGT events, others may come from technical artifacts such as bacterial contamination of eukaryotic sequences. To minimize such false positive HGT calls, we took a conservative approach in our analyses, considering potential bacteria-eukaryote HGT events to be trustworthy only if: 1) eukaryotic and bacterial sequences branched near one another with strong support (bootstrap values >70); 2) the eukaryotic sequences formed a distinct subclade, represented by at least 2 species from the same eukaryotic supergroup; 3) the eukaryotic sequences were produced by at least 2 different studies; and 4) the position of the horizontally transferred sequences was robust across all alignment and phylogenetic reconstruction methods used (Supp. Fig. 3). While these restrictions limit our attention to relatively old HGT events, they also give us confidence these events are likely to be real.

The Mab21 superfamilies passed all four of these HGT thresholds, as did another eukaryotic clade of CD-NTases that were all previously undescribed. We name this clade the eukaryotic SMODS (eSMODS) superfamily, because the top scoring domain from hmmscan for each sequence in this superfamily was the SMODS domain (PF18144), which is typically found only in bacterial CD-NTases (Supplementary Data). This sequence similarity suggests that eSMODS arose following a recent HGT from bacteria and/or that these CD-NTases have diverged from their bacterial predecessors less than the eukaryotic OAS and Mab21 families have. Additionally, all of these sequences were predicted to have a Nucleotidyltransferase domain (PF01909), and (8/12) had a Polymerase Beta domain (PF18765), which are features shared with many bacterial CD-NTases in Clades D,E, and F (Supplementary Data). The eSMODS superfamily is made up of sequences from Amoebozoa, choanoflagellates, Ancryomonadida, and one animal (the sponge *Oscarella pearsei*), which clustered robustly and with high support within bacterial Clade D(e.g. subclade D04, CD-NTase 22 from *Myxococcus xanthus*) (Supp. Fig 4). The eSMODS placement on the tree, which was robust to all alignment and phylogenetic algorithms used (Supp. Fig. 3), suggesting that eSMODS represent an additional, independent acquisition of CD-NTases from bacteria.

CD-NTases from bacterial Clade C and Clade D are the only CD-NTases to produce cyclic trinucleotides, producing cyclic tri-Adenylate and cAAG, respectively[11,47–49]. Interestingly, OAS produces linear adenylates, which is one step away from the cAAA product made by Class C CD-NTases, and similarly cGAMP (made by cGAS) is one adenylate away from the class D product cAAG. As of this writing, the Clade D CD-NTases closest to the eSMODS and Mab21 superfamilies (D04 and D12, respectively), have not been well characterized. Therefore we argue that these CD-NTases should be a focus of future studies, as they may hint at the evolutionary stepping stones that allow eukaryotes to acquire bacterial immune proteins.

### Diverged eukaryotic STINGs bridge the gap between bacteria and animals

We next turned to analyze Stimulator of Interferon Gamma (STING) proteins. In animals, STING is a critical cyclic dinucleotide sensor, important during viral, bacterial, and parasitic infections (reviewed here[50]). Structurally, most metazoan STINGs consist of an N-terminal transmembrane domain (TM), made of 4 alpha helices fused to a C-terminal STING domain[51]. Canonical animal STINGs show distant homology with STING effectors from the bacterial cyclic oligonucleotide-based antiphage signaling system (CBASS), with major differences in protein structure and pathway function between these animal and bacterial defenses. For example, in bacteria, the majority of STING proteins are fusions of a STING domain to a TIR (Toll/interleukin-1 receptor) domain (Fig. 3A). Bacterial STING proteins recognize cyclic di-GMP and oligomerize upon activation, which promotes TIR enzymatic activity[3,52,53]. Some bacteria, such as *Flavobacteriaceae,* encode proteins that fuse a STING domain to a transmembrane domain, although it is unclear how these bacterial TM-STINGs function[3]. Other bacteria have STING domain fusions with deoxyribohydrolase, ⍺/β- hydrolase, or trypsin peptidase domains[14]. In addition to eukaryotic TM-STINGs, a few eukaryotes such as the oyster *Crassostrea gigas* have TIR-STING fusion proteins, although the exact role of their TIR domain remains unclear[3,54,55].

**Figure 3:**
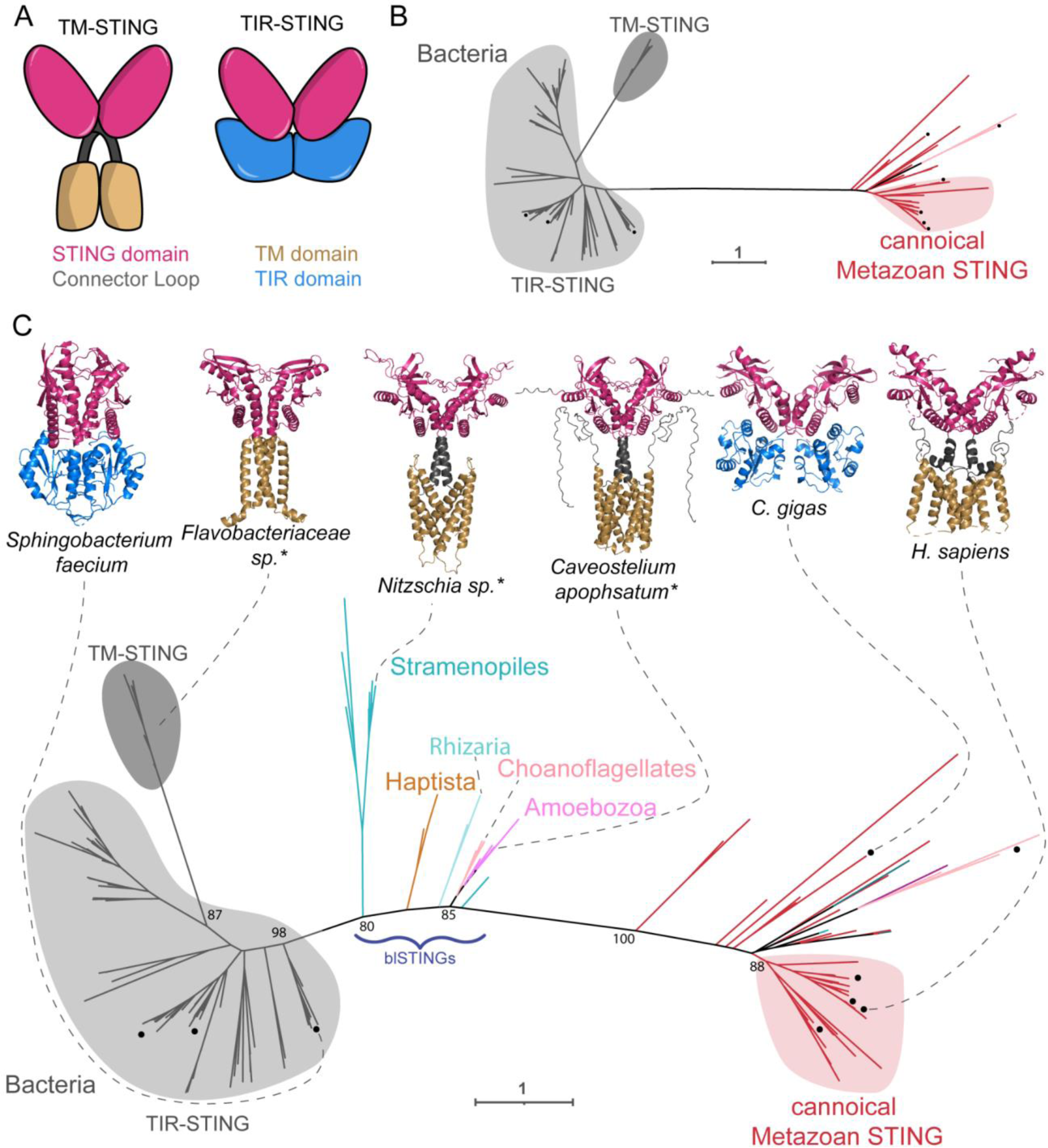
Diverse eukaryotic STING proteins bridge the gap between metazoans and bacteria. (A) Graphical depiction of common domain architectures of STING proteins. (B) Maximum likelihood unrooted phylogenetic tree of STING domains from Metazoa and bacteria, which are separated by one long branch. Black dot (•) indicates proteins that have been previously experimentally characterized. Bacterial sequences are in gray and animal sequences are in red. (C) Maximum likelihood unrooted phylogenetic tree of hits from iterative HMM searches for diverse eukaryotic STING domains. The STING domains from bacteria-like STINGs (blSTINGs) from diverse eukaryotes break up the long branch between bacterial and animal STINGs. Structures of the indicated STING proteins are shown above, with those predicted by AlphaFold indicated by an asterisk. Homologs with X-ray crystal structures are from[3,82]. Colored regions show two domain architectures in bacteria and eukaryotes (STING linked to a TIR domain and STING linked to a transmembrane domain), each of which have evolved convergently in bacteria and eukaryotes. Ultrafast bootstraps determined by IQtree shown at key nodes. See Supplementary Figure 5 for full STING phylogenetic tree.

Given these major differences in domain architectures, ligands, and downstream immune responses, how have animals and bacteria evolved their STING-based defenses, and what are the relationships between them? Prior to this work, the phylogenetic relationship between animal and bacterial STINGs has been difficult to characterize with high support[14]. Indeed, when we made a tree of previously known animal and bacterial STING domains, we found that the metazoan sequences were separated from the bacterial sequences by one very long branch, along which many changes had occurred (Fig. 3B).

To improve the phylogeny through the inclusion of a greater diversity of eukaryotic STING sequences, we began by carefully identifying the region of STING that was homologous between bacterial and animal STINGs, as we expected this region to be best conserved across diverse eukaryotes. Although Pfam domain PF15009 (TMEM173) is commonly used to define animal STING domains, this HMM includes a portion of STING’s transmembrane domain which is not shared by bacterial STINGs. Therefore, we compared the crystal structures of HsSTING (6NT5), *Flavobacteriaceae sp.* STING (6WT4) and *Crassostrea gigas* STING (6WT7) to define a core “STING” domain. We used the region corresponding to residues 145-353 of 6NT5 as an initial HMM seed alignment of 15 STING sequences from PF1500915 (“Reviewed” sequences on InterPro). Our searches yielded 146 eukaryotic sequences from 64 species, which included STING homologs from 34 metazoans (Supplemental Data and Fig. 1). Using maximum likelihood phylogenetic reconstruction, we identified STING-like sequences from 26 diverse microeukaryotes that clustered in between bacterial and metazoan sequences, breaking up the long branch. We name these sequences the bacteria-like STINGs (blSTINGs) because they were the only eukaryotic group of STINGs with a bacteria-like Prok_STING domain (PF20300) and due to the short branch length (0.86 vs. 1.8) separating them from bacterial STINGs on the tree (Fig. 3C). While a previous study reported STING domains in two eukaryotic species (one in Stramenopiles and one in Haptista) [14], we were able to expand this set to additional species and also recover blSTINGs from Amoebozoa, Rhizaria and choanoflagellates. This diversity allowed us to place the sequences on the tree with high confidence, recovering a substantially different tree than previous work[14]. As for CD-NTases, the tree topology we recovered was robust across multiple different alignment and phylogenetic tree construction algorithms (Supp. Fig. 3).

Given the similarities between the STING domains of the blSTINGs and bacterial STINGs, we next asked whether the domain architectures of these proteins were similar using Hmmscan and AlphaFold. The majority of the new eukaryotic blSTINGs were predicted to have four N-terminal alpha helices (Fig. 3A, and Supplementary Data), similar to human STING. While bacterial TM-STINGs were superficially similar with N-terminal transmembrane domains, these proteins were predicted to have only two alpha helices and in 5/6 phylogenetic trees bacterial TM-STINGs were more similar to other bacterial STINGs than to eukaryotic homologs (Supp. Fig. 3). These results suggest that eukaryotes and bacteria independently converged on a common TM-STING domain architecture through domain shuffling.

Interestingly, a similar pattern of convergent domain shuffling appears to have occurred a second time with the TIR-STING proteins. It was previously known that some eukaryotes such as the oyster *C. gigas,* have a TIR-STING fusion protein[3,54,55]. The STING domain of these TIR-STINGs clustered closely to other metazoan STINGs, suggesting an animal origin (Fig. 3B). We also investigated the possibility that *C. gigas* acquired the TIR-domain of its TIR-STING protein via HGT from bacteria, however this analysis also suggested an animal origin for the TIR domain (Supp. Fig. 7). Eukaryotic TIR-STINGs are rare, further supporting the hypothesis that this protein resulted from recent convergence, where animals independently fused STING and TIR domains to make a protein resembling bacterial TIR-STINGs, consistent with previous reports[14]. Overall, we find that the TM-STING and TIR-STING proteins represent at least two independent examples of convergent evolution, where bacteria and eukaryotes have created similar proteins through the reuse of ancient protein domains. Our work also identified a number of non-metazoan STINGs (the blSTINGs) that have a domain architecture similar to animal STINGs but a STING domain more similar to bacterial STINGs.

### Viperin is an ancient and widespread immune family

Viperins are innate immune proteins that restrict the replication of a diverse array of viruses by conversion of nucleotides into 3’-deoxy-3’,4’didehdro-(ddh) nucleotides[4,56–58]. Incorporation of these ddh nucleotides into a nascent RNA molecule leads to chain termination, blocking RNA synthesis and inhibiting viral replication[56,59]. While metazoan viperin specifically catalyzes CTP to ddhCTP[56], homologs from archaea and bacteria can generate ddhCTP, ddhGTP, and ddhUTP[4,60]. Previous structural and phylogenetic analysis showed that eukaryotic viperins are highly conserved at both the sequence and structural level and that, phylogenetically, animal and fungal viperins form a distinct monophyletic clade compared to bacterial viperins[4,57,60].

As viperin proteins consist of a single Radical SAM protein domain, we iteratively searched EukProt beginning with domain PF04055 (Radical_SAM). The 194 viperin-like proteins we recovered came from 158 species spanning the full range of eukaryotic diversity, including organisms from all of the major eukaryotic supergroups, as well as some orphan taxa whose taxonomy remains open to debate (Fig. 1). When we constructed phylogenetic trees from these sequences, we found that the large majority of the eukaryotic viperins cluster together in a single, monophyletic clade, separate from bacterial or archaeal viperins (Fig. 4). Within the eukaryotic viperin clade, sequences from more closely related eukaryotes often clustered together (Fig. 4, colored blocks), as would be expected if viperins were present and vertically inherited within eukaryotes for an extended period of time. The vast species diversity and tree topology both strongly support the inference that viperins are a truly ancient immune module and have been present within the eukaryotic lineage likely dating back to the last eukaryotic common ancestor (LECA).

**Figure 4:**
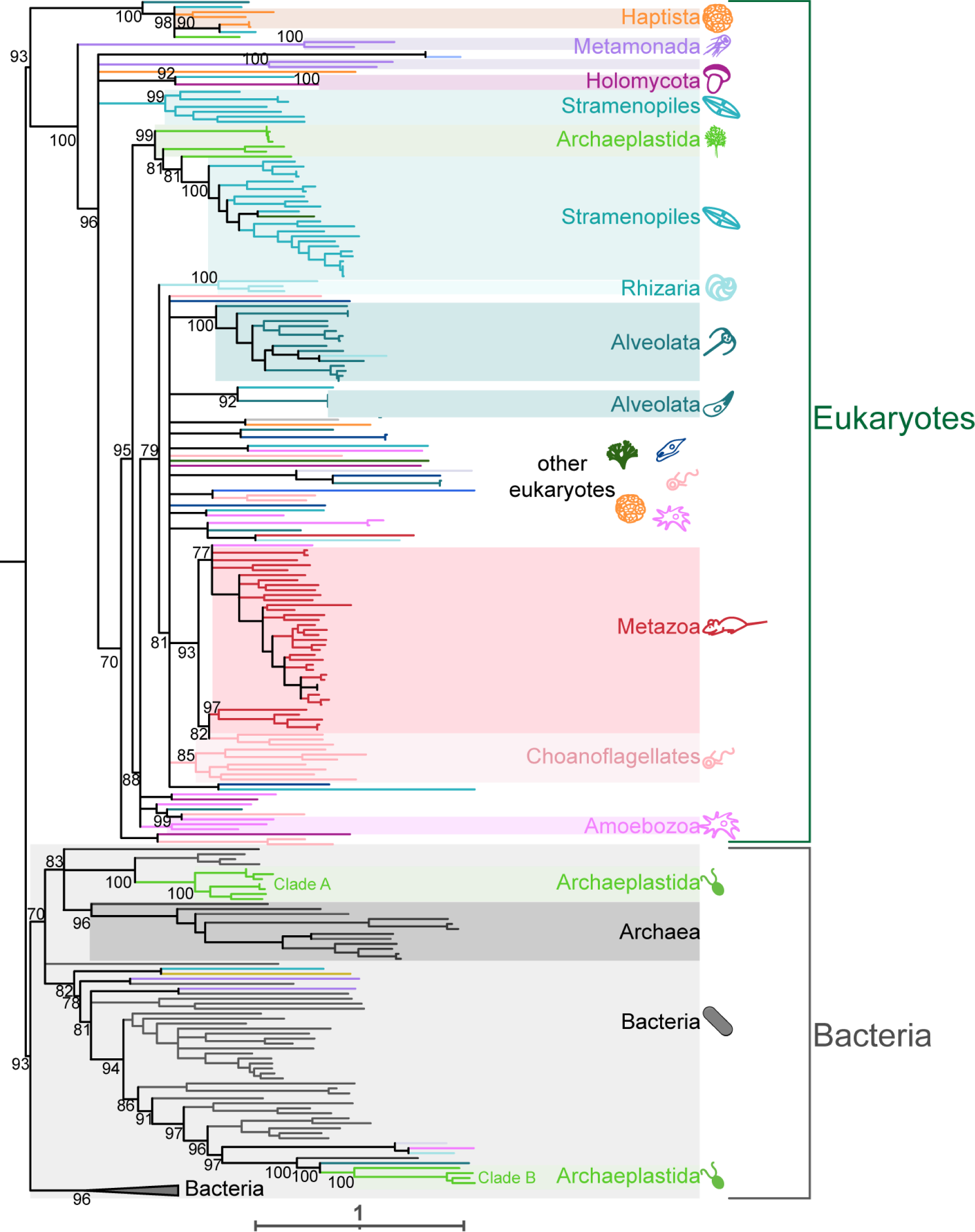
Viperin is a deeply conserved innate immune module. (A) Maximum likelihood phylogenetic tree generated by IQtree of Viperins from eukaryotes, bacteria, and archaea. All major eukaryotic supergroups have at least two species that encode a Viperin homolog (colored supergroups). Bacterial Viperin sequences shown in gray and archaeal sequences in dark gray. There are two clades of Chloroplastida (a group within Archaeplastida) sequences that branch robustly within the bacteria clade. Ultrafast bootstraps determined by IQtree shown at key nodes. Tree is arbitrarily rooted between the major eukaryotic and bacterial clades. See Supplementary Figure 6 for fully annotated Viperin phylogenetic tree.

In addition to this deep eukaryotic ancestry, we also uncovered two examples of bacteria-eukaryote HGT that have occurred much more recently, both in Chloroplastida, a group within Archaeplastida. The first of these consists of a small clade of Archaeplastida (Clade A) consisting of marine algae such as *Chloroclados australicus* and *Nemeris dumetosa*. These algal viperins cluster closely with the marine cyanobacteria *Anabaena cylindrica* and *Plankthriodies* (Fig. 4 and Supp. Fig. 6). The second clade (Clade B) includes four other Archaeplastida green algal species, mostly *Chlamydomonas spp*. In some of our trees the Clade B viperins branched near to eukaryotic sequences from other eukaryotic supergroups, however the placement of the neighboring eukaryotic sequences varied depending on the algorithms we used; only the Archaeplastida placement was consistent. (Fig. 4 and Supp. Fig. 3 & 6). Taken together, we conclude that viperins represent a class of ancient immune proteins that have likely been present in eukaryotes since the LECA. Yet, we also find ongoing evolutionary innovation in viperins via HGT, both among eukaryotes and between eukaryotes and bacteria.

## Discussion

The recent discoveries that bacteria and mammals share mechanisms of innate immunity have been surprising, because they imply that there are similarities in immunity that span the Tree of Life. But how did these similarities come to exist? Here we uncover several evolutionary trajectories that have led animals and bacteria to share homologous immune proteins (summarized in Fig. 5). We found that Viperin is truly ancestral, dating back to at least the Last Eukaryotic Common Ancestor (LECA), and likely further. We also uncovered examples of convergence, as in STING, where the shuffling of ancient domains has led animals and bacteria to independently arrive at similar protein architectures. Finally, we found evidence of multiple examples of bacteria-eukaryote HGTs that have given rise to immune protein families. An essential part of our ability to make these discoveries was the analysis of data from nearly 1000 diverse eukaryotic taxa. These organisms allowed us to distinguish between proteins found across eukaryotes vs. animal-specific innovations, to document both recent and ancient HGT events from bacteria that gave rise to eukaryotic immune protein families (Fig. 2 & 4), and to identify STING proteins with eukaryotic domain architectures but more bacteria-like domains (blSTINGs, Fig. 3). Because these diverged eukaryotic STINGs were found in organisms where we typically did not find any CD-NTase proteins, we hypothesize that blSTINGs may detect and respond to exogenous cyclic nucleotides, such as those generated by pathogens. In contrast to the STINGs, the eukaryotic CD-NTases had substantially different evolutionary histories, with multiple major CD-NTase superfamilies each emerging from within larger bacterial clades. While these analyses cannot definitively determine the directionality of the transfer, we favor the most parsimonious explanation that these components came into the eukaryotic lineage from bacterial origins.

**Figure 5:**
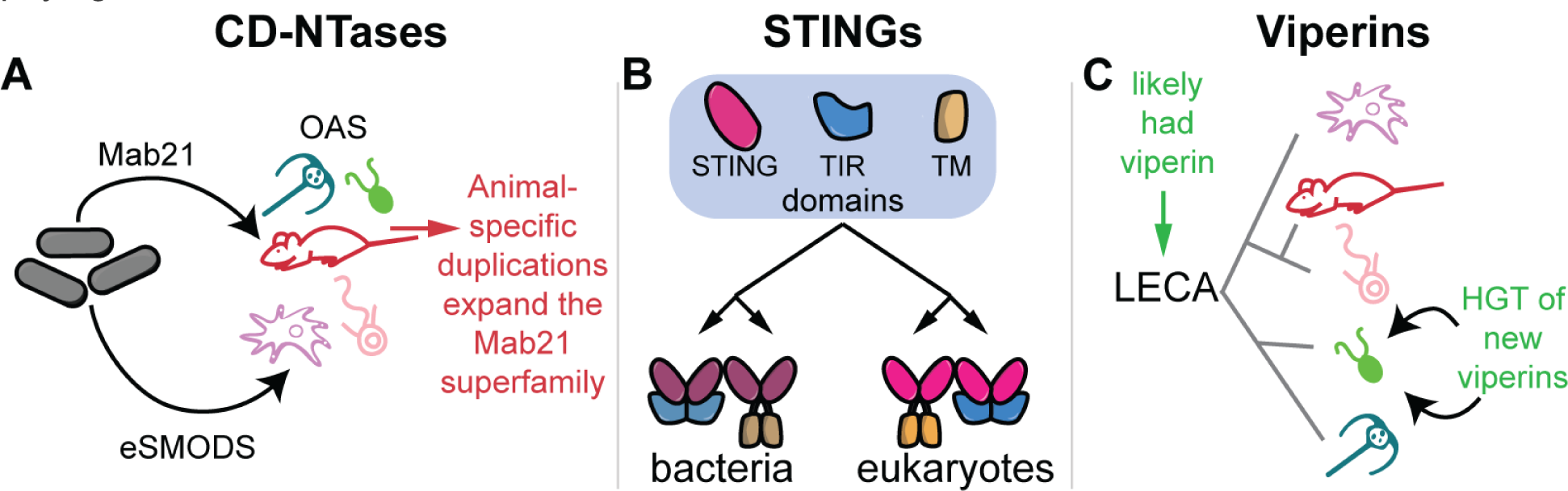
Proposed model of evolutionary history of CD-NTases, STING, and Viperin. Proposed summary models of the evolutionary history of innate immune components. (A) We define two distinct superfamilies of CD-NTases that likely arose from bacteria-eukaryote HGT: eSMODS and Mab21. Within the Mab21 superfamily (which contains cGAS), a number of animal-specific duplications gave rise to numerous paralogs. The OAS superfamily of CD-NTases are abundant across diverse eukaryotic taxa and were likely present in the LECA. (B) Drawing on a shared ancient repertoire of protein domains that includes STING, TIR, and transmembrane (TM) domains, bacteria and eukaryotes have convergently evolved similar STING proteins through domain shuffling. (C) Viperins are widespread across the eukaryotic tree and likely were present in the LECA. In addition, two sets of recent HGT events from bacteria have equipped algal species with new viperins.

While not as prevalent as in bacteria, HGT in eukaryotes represents a significant force in eukaryotic evolution, especially for unicellular eukaryotes[61–64]. In this study, our criteria for ‘calling’ HGT events was relatively strict, meaning that our estimate of HGT events is almost certainly an underestimate. Importantly, this pattern suggests that the bacterial pan-genome has been a rich reservoir that eukaryotes have repeatedly sampled to acquire novel innate immune components. Some of these HGT events have given rise to new eukaryotic superfamilies (e.g. eSMODS) that have never been characterized and could represent novel types of eukaryotic immune proteins. We speculate that the eSMODS superfamily CD-NTases and the blSTINGs may function more similarly to their bacterial homologs, potentially producing and responding to a variety of cyclic di- or tri-nucleotides[11] Similarly, bacterial viperins have been shown to generate ddhCTP, ddhGTP, and ddhUTP, whereas animal viperins only make ddhCTP[4,56,60]. Thus, the two algal viperin clades arising from HGT may have expanded functional capabilities as well. A caveat of this work is that such strictly bioinformatic investigations are insufficient to reveal protein biochemical functions, nor can they determine whether diverse homologs have been co-opted for non-immune functions. We therefore urge future, functional studies to focus on these proteins to resolve the questions of 1) whether/how blSTINGs operate in the absence of CD-NTases, 2) whether/how the functions of algal viperins and eSMODS changed following their acquisition from bacteria, and 3) whether the homologs truly function in immune defense.

In addition to these instances of gene gain, eukaryotic gene repertoires have been dramatically shaped by losses. Even for viperins, which likely date back to the eukaryotic last common ancestor, these proteins were sparsely distributed across eukaryotes and were absent from the majority of species we surveyed. While some of this finding may be due to technical limitations, such as dataset incompleteness or inability of the HMMs to recover distant homologs, we believe this explanation is insufficient to fully explain the sparseness, as many plant, fungal, and amoebozoan species are represented by well-assembled genomes where these proteins are certifiably absent (Supp. Fig. 2). Instead, we propose that the sparse distribution likely arises from ongoing and repeated gene loss, as has been previously documented for other gene families across the Tree of Life[22,24–26].

Overall, our results yield a highly dynamic picture of immune protein evolution across eukaryotes, wherein multiple mechanisms of gene gain are offset by ongoing losses. Interestingly, this pattern mirrors the sparse distributions of many of these immune homologs across bacteria[65–67], as anti-phage proteins tend to be rapidly gained and lost from genomic defense islands[68,69]. It will be interesting to see if some eukaryotes evolve their immune genes in similarly dynamic islands, particularly in unicellular eukaryotes that undergo more frequent HGT[70].

We expect that our examination of STING, CD-NTases, and Viperin represents just the tip of the iceberg when it comes to the evolution of eukaryotic innate immunity. New links between bacterial and animal immunity continue to be discovered and other immune families and domains such as Argonaute, Gasdermins, NACHT domains, CARD domains, TIR domains, and SamHD1 have been shown to have bacterial roots[2,6,7,9,10]. To date, the majority of studies have focused on proteins specifically shared between metazoans and bacteria. We speculate that there are probably many other immune components shared between bacteria and eukaryotes outside of animals. Further studies of immune defenses in microeukaryotes are likely to uncover new mechanisms of cellular defense and to better illustrate the origins and evolution of eukaryotic innate immunity.

## Methods

### Iterative HMM Search

The goal of this work was to search the breadth of EukProt v3 for immune proteins from the CD-NTase, STING, and viperin families that span the gap between metazoan and bacterial immunity. Our overall strategy was to first search with eukaryotes alone (starting from mainly Metazoa). Then we added in bacterial sequences and searched with a mixed bacterial-eukaryotic HMM search until we either found no new hits, or until we began getting hits from an outgroup gene family. In parallel, we also performed bacteria-only and eukaryote-only searches, to ensure that we found as many homologs as possible (schematized in Fig 1A, and further in Supp. Fig. 1A).

#### Phase 1: Eukaryotic searches

To begin, hidden Markov model (HMM) profiles from Pfam (for CD-NTases and Viperin) or an HMM profile generated from a multiple sequence alignment (for STING) were used to search EukProt V3[15], for diverse eukaryotic sequences. For CD-NTases and Viperin, HMM profiles of Pfams PF03281 and PF04055 were used respectively.

For STING, where the Pfam profile includes regions of the protein outside of the STING domain, we generated a new HMM for the initial search. First, we aligned crystal structures of HsSTING (6NT5), *Flavobacteriaceae sp.* STING (6WT4) and *Crassostrea gigas* STING (6WT7) to define a core “STING” domain. Then we aligned 15 eukaryotic sequences from PF15009 (“Reviewed” sequences on InterPro) with MAFFT(v7.4.71)[71] and manually trimmed the sequences down to the boundaries defined by our crystal alignment (residues 145-353 of 6NT5). We then trimmed the alignment with TrimAI (v1.2)[72] with options -gt 0.2. The trimmed MSA was then used to generate an HMM profile with hmmbuild from the hmmer (v3.2.1) package (hmmer.org).

HMM profiles were used to search EukProt via hmmsearch (also from hmmer v3.2.1) with a statistical cutoff value of 1e-3 and -hit parameter set to 10 (i.e. the contribution of a single species to the output list is capped at 10 sequences). The resulting hits from this search were then aligned with hmmalign (included within hmmer) and used to generate a new HMM profile with hmmbuild. This profile was used to search EukProt v3 again and the process was repeated for a total of 3-4 eukaryotic searches.

#### Phase 2: combining eukaryotic and bacterial sequences into an HMM

After the eukaryotic searches reached saturation (i.e. no additional eukaryotic sequences were recovered after additional searches), bacterial sequences were acquired from previous literature (Viperins from[4], CD-NTases from[11], and STINGs from[3,8,43]). To ensure the combined HMM did not have an overrepresentation of either bacterial or eukaryotic sequences, we downsampled the bacterial sequences and eukaryotic sequences to obtain 50 phylogenetically diverse sequences of each, and then combined the two downsampled lists. To do this, eukaryotic and bacterial sequences were each separately aligned with MAFFT, phylogenetic trees were built with FastTree (v2.1.10)[73], and the Phylogenetic Diversity Analyzer (pda/1.0.3)[74] software was used to downsample the sequences while maximizing remaining sequence diversity.

The combined bacterial-eukaryotic sequence list was then aligned with hmmalign and used to construct a new HMM profile with hmmbuild. This HMM profile was used to search EukProt v3. The eukaryotic hits from this search were then aligned with MAFFT, and a tree was constructed with FastTree. From this tree the sequences were then downsampled with PDA and once again combined with the bacterial list, aligned, used to generate a new HMM, and a new search. This process was iterated 3-5 times until saturation or until the resulting sequence hits included other gene families that branched outside of the sequence diversity defined by the metazoan and bacterial homologs.

#### Phase 3: Searching with a bacteria-only or existing eukaryote-only HMM profiles

To search EukProt v3 with a bacteria-only HMM for each protein family, we aligned the full set of published bacterial sequences with MAFFT, trimmed with TrimAI, and hmmbuild was used to generate an HMM profile which was used to search EukProt v3. As a point of comparison, we also searched the database with only the starting, animal-dominated Pfam (PF15009) for STING.

#### Phase 4: Combining all hits into a single list and scanning for domains

Sequences from all iterative searches were combined to generate a total hits FASTA file for STING, CD-NTase, and Viperin. First, duplicate sequences were removed, then the fasta files were scanned using hmmscan (also from hmmer v3.2.1) against the Pfam database (Pfam-A.hmm) and all predicted domains with an E-value <1e-3 were considered. Next, we generated phylogenetic trees (first by aligning with MAFFT and then building a tree with FastTree), and used these trees along with the hmmscan domains to determine in-group and out-group sequences. Out-group sequences were removed from the fasta file. We determined outgroup sequences by these criteria: 1) if the sequence clustered outside of known outgroup sequences (e.g. Poly(A) RNA polymerase (PAP) sequences for the CD-NTases, and molybdenum cofactor biosynthetic enzyme (MoaA) for Viperin), or 2) if sequence did not have at least one of the relevant domains (Mab21/OAS1-C/SMODS for CD-NTases, TMEM173/Prok_STING for STING, and Radical_SAM for Viperin). These three FASTA files were used for the final alignments and phylogenetic trees. To identify protein domains in each sequence, the FASTA files were scanned using hmmscan (also from hmmer v3.2.1) against the Pfam database (Pfam-A.hmm) and all predicted domains with an E-value <1e-3 were considered. See Supplemental Data for the hmmscan results of all included homologs.

### Final Alignment and Tree Building

To generate final phylogenetic trees, all eukaryotic search hits and bacterial sequences were aligned using MAFFT. We downsampled the CD-NTase bacterial sequences from ∼6000 down to 500 as described above, to facilitate more manageable computation times on alignments and tree construction. For the STING and Viperin trees we included all bacterial sequences. These initial alignments were first trimmed manually in Geneious (v2023.1.2) to remove unaligned N- and C-terminal regions, and then realigned with MAFFT or MUSCLE(v5.1)[75] and trimmed with TrimAI (v1.2)[72]. MUSCLE was used with the “-super5” option, and otherwise default parameters. These alignments were used to generate phylogenetic trees using three tree inference softwares: FastTree (v2.1.10)[73], IQtree (2.2.2.7)[76] and RaxML-ng (v0.9.0)[77]. FastTree was utilized with default settings. IQtree was used to determine the appropriate evolutionary model, and was run with 1000 ultrafast bootstraps (IQtree settings: -s, -bb 1000, -m TEST, -nt AUTO). RaxML-ng trees were produced with 100 bootstraps using the molecular model specified from the IQtree analysis (Raxml-ng settings: --all, --model [specified by IQtree], --tree pars{10} --bs-trees 100). Phylogenetic trees were visualized with iTOL[78].

### TIR Domain Alignment and Tree

We used hmmscan to identify the coordinates of TIR domains in a list of 203 TIR domain containing-sequences from InterPro (Family: IPR015032) and 104 bacterial TIR-STING proteins (the same TIR-STING proteins used in Fig. 3)[3]. Next, we trimmed the sequences down to the TIR coordinates and aligned the TIR domains with MUSCLE. We trimmed the alignments with TrimAL and built a phylogenetic tree with IQtree.

### Venn Diagrams

Venn diagrams were generated via DeepVenn[79] using presence/absence information for Mab21, OAS, and STING from each eukaryotic species that encoded at least one of these proteins.

### Protein Structure Modeling

To model 3-D protein structures for STING homologs without a published crystal structure, we ran AlphaFold (v2.1.1)[80,81]. We generated 5 ranked models for STINGs from *Flavobacteriaceae* (IMG ID: 2624319773), *Nitzschia sp.* (EukProt ID: P007051), and *Caveostelium apophsatum* (EukProt ID: P019191). Fig. 2C shows highest ranked models only.

## Acknowledgements

We thank Daniel Richter for his feedback, encouragement, and scientific guidance. Maureen Stolzer, Kevin Forsberg, Patrick Mitchell, and members of the Levin lab also provided helpful input on the project and manuscript. Thanks to Andrew VanDemark for helpful discussions about 3-D modeling and to Jacob Durrant for help running AlphaFold. This research was supported in part by the University of Pittsburgh Center for Research Computing, RRID:SCR_022735, through the resources provided. Specifically, this work used the HTC cluster, which is supported by NIH award number S10OD028483. Ed Culbertson was supported by NSF Postdoctoral fellowship 2208971 and Tera Levin was supported by NIH R00AI139344 and R35GM150681.

---

**Supplementary Figure 1:**
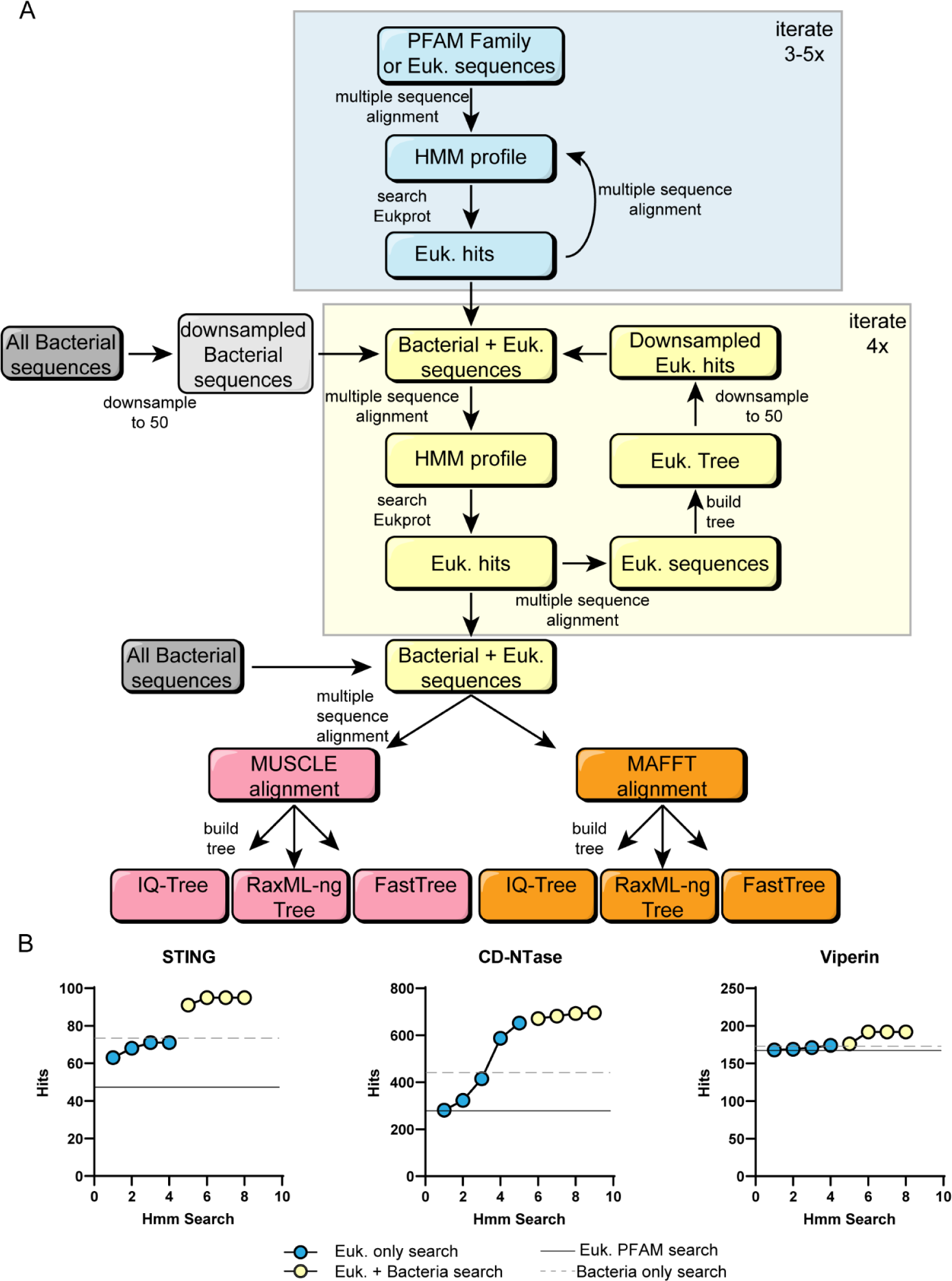
Collectors curves and full search strategy. (A) Detailed schematic outlining the iterative HMM search strategy. Blue boxes and blue shaded region show eukaryotic-only searches to create pan-eukaryotic HMMs and yellow indicates eukaryotic-bacterial searches to create universal HMMs. For the combined bacterial/eukaryotic searches (yellow box), bacterial and eukaryotic sequences were each downsampled to 50 sequences (phylogenetic tree downsampled via PDA) to maintain equal contributions from bacteria and eukaryotic sequences. Separately, bacterial sequences were aligned and used to make an HMM which was used to search EukProt as a ‘bacteria only search’ and for STING we searched with PF15009 for a comparable Eukaryotic PFAM search (not shown in flowchart). We did this extra search for STING as PF15009 contains part of the eukaryotic STING transmembrane domain and so our first search with STING was with a STING-domain-only HMM (See Materials and Methods). Pink (MUSCLE) and orange (MAFFT) boxes show the final alignments and phylogenetic trees that were constructed. (B) STING, CD-NTase, and Viperin collector’s curves showing the number of cumulative protein sequences that were found after each iterative search. Results from eukaryotic searches are shown in blue and the combined searches in yellow. Solid black line indicates the number of hits from the starting Pfam HMM alone and the dotted gray line shows the number of hits from a bacteria-only HMM. Note that some searches yielded hits that were members of more distant protein families, which were later removed from the analysis and are not counted here.

**Supplementary Figure 2:**
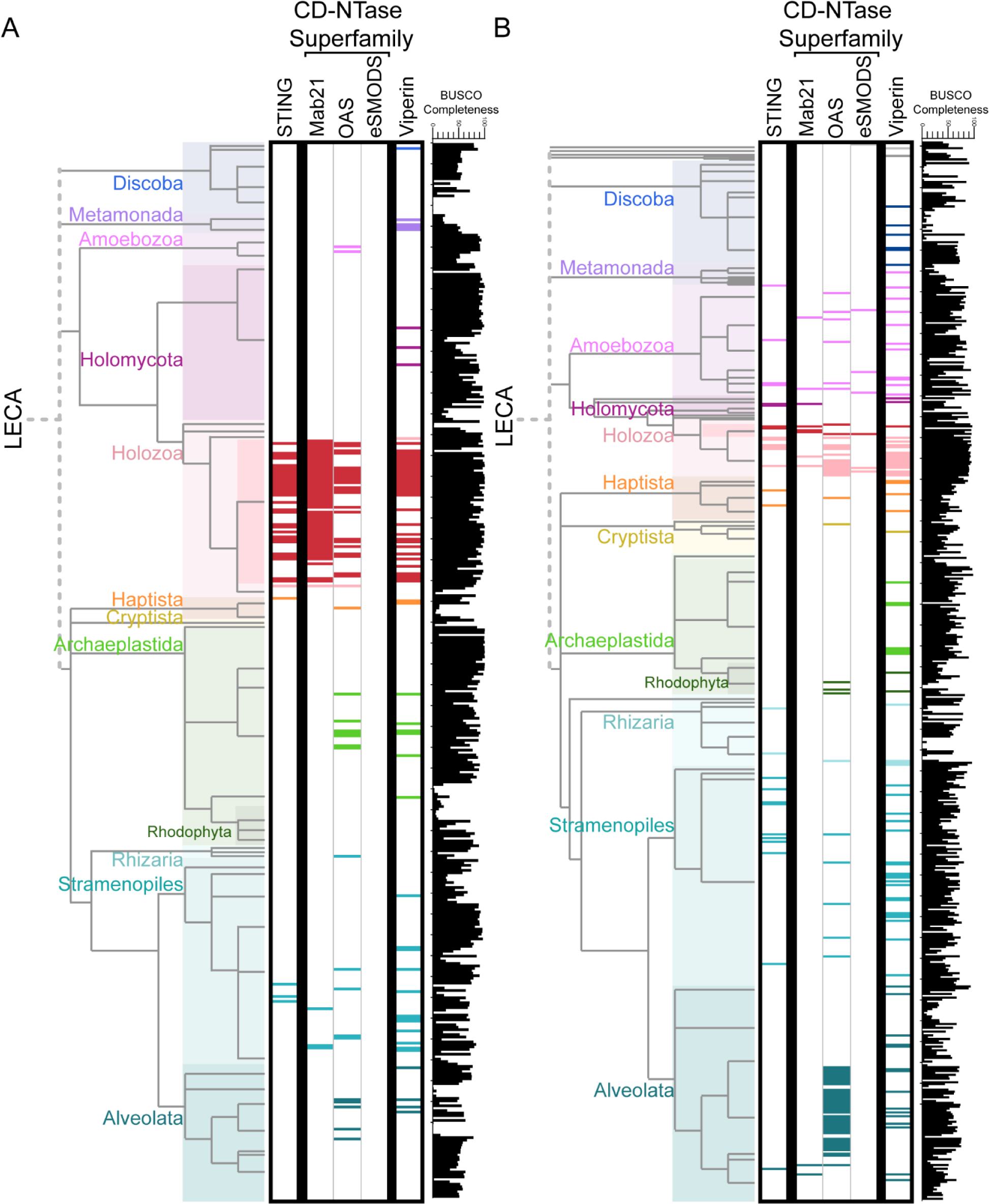
Phylogenetic trees of EukProt species by data type. Phylogenetic trees derived from Fig. 1 separating species represented in EukProt v3 by genomes (A) or transcriptomes (B). Supergroups are color-coded as in Figure 1. Colored bars mark each eukaryotic species in which the HMM search found a homolog sequence of STING, CD-NTase, or Viperin. Black bar chart shows BUSCO completeness score for each genome/transcriptome.

**Supplementary Figure 3:**
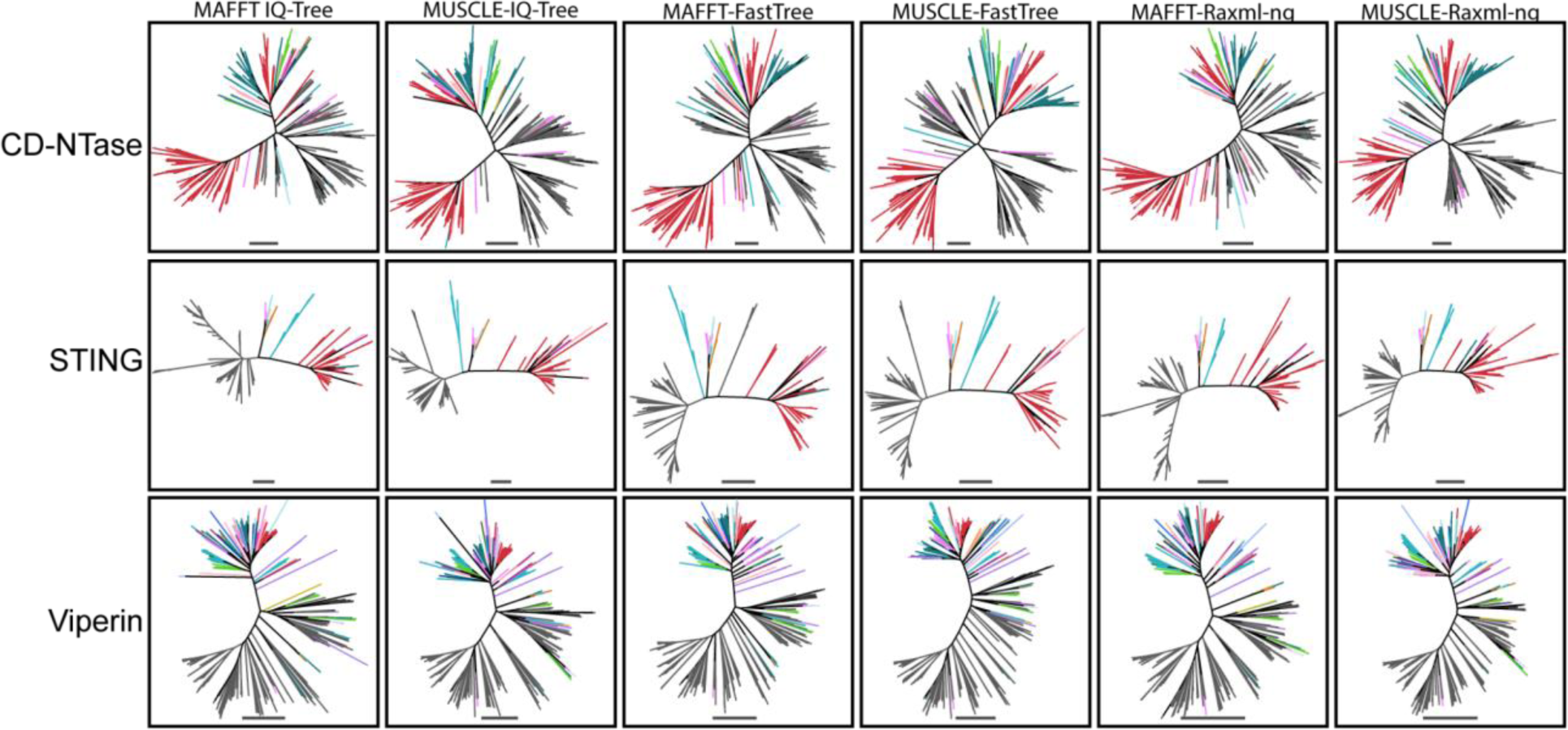
Phylogenetic trees from different alignments and tree building methods show robust topologies. Unrooted maximum likelihood phylogenetic trees generated from two separate alignments (MUSCLE and MAFFT) and with three different tree inference programs (FastTree, IQtree, and RaxML-ng). Scale bar of 1 shown beneath each tree represents the number of amino acid substitutions per position in the underlying alignment. Colored branches show eukaryotic sequences with the same color scheme as Fig. 1, while gray lines are bacterial sequences. For the majority of relationships discussed here, we recovered the same tree topology at key nodes regardless of alignment or tree reconstruction algorithm used. The only exception was in the STING FastTree phylogenies, wherein the TM-STING clade moved to multiple positions in the phylogeny, depending on alignment algorithm used.

**Supplementary Figure 4:**
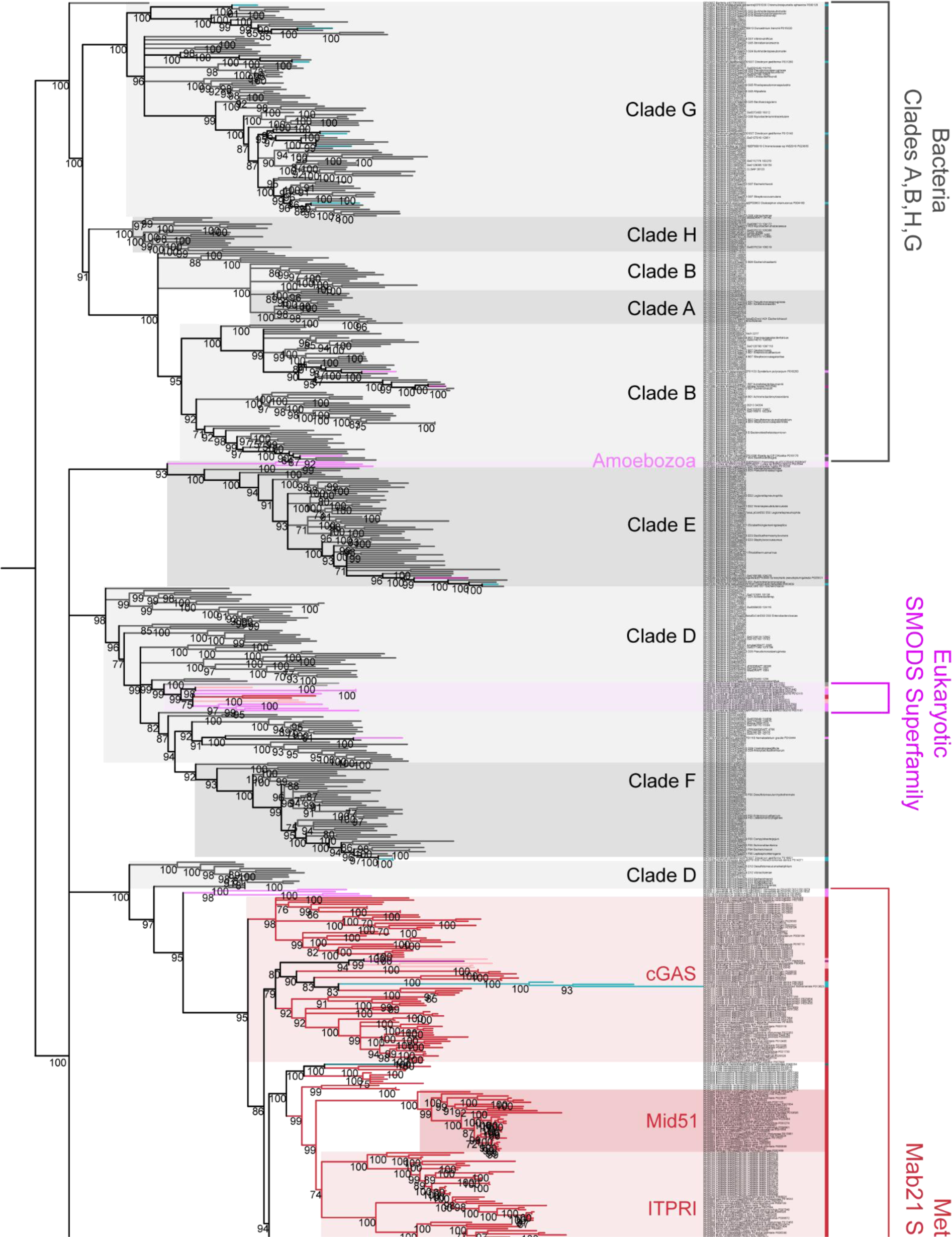
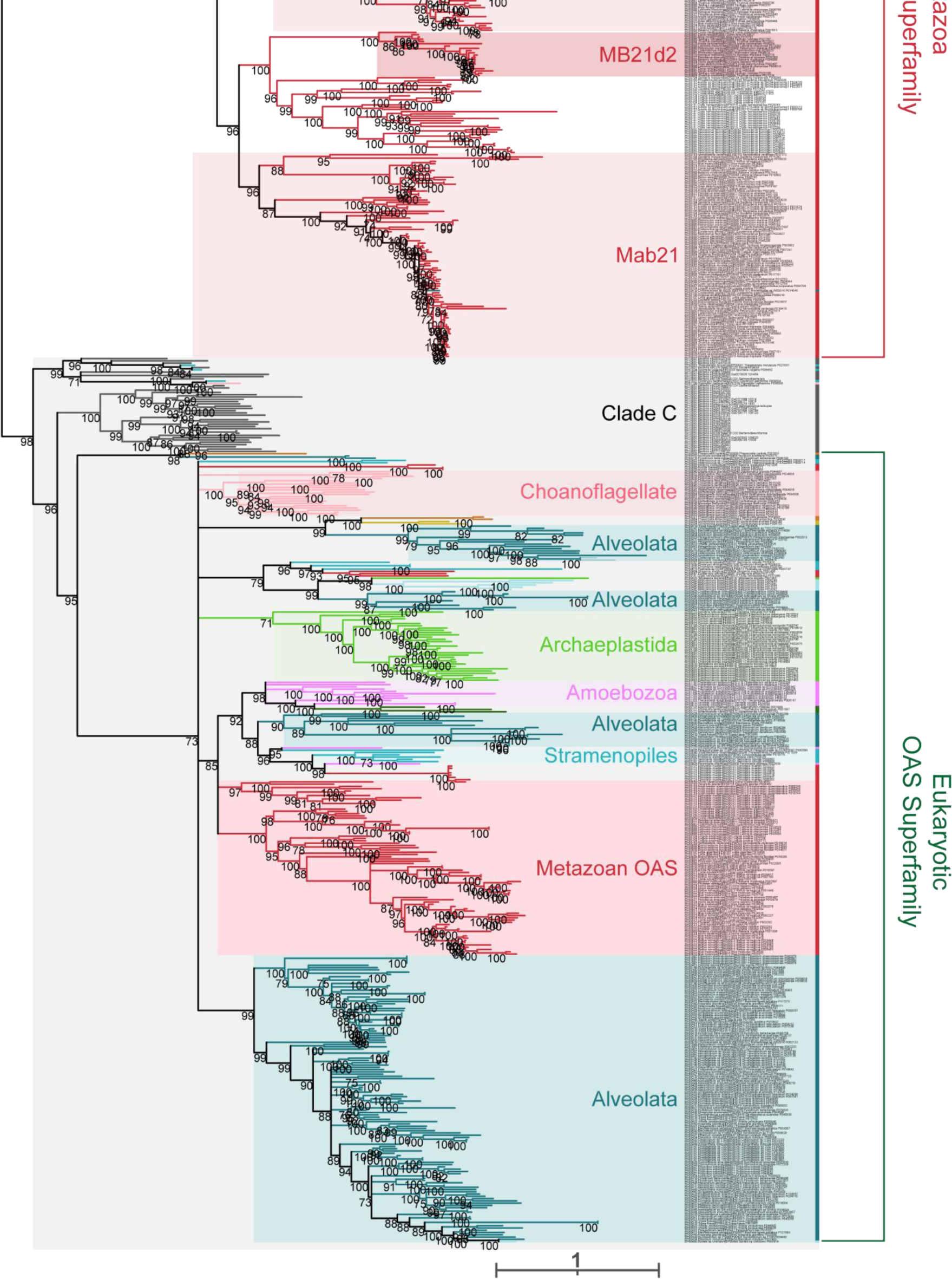
CD-NTase phylogenetic tree. Maximum likelihood phylogenetic tree generated by IQtree of hits from iterative HMM searches for diverse eukaryotic CD-NTases. Scale bar represents the number of amino acid substitutions per position in the underlying MUSCLE alignment. Ultrafast bootstrap values calculated by IQtree at all nodes with support >70 are shown. Branches with support values <70 were collapsed to polytomies.

**Supplementary Figure 5:**
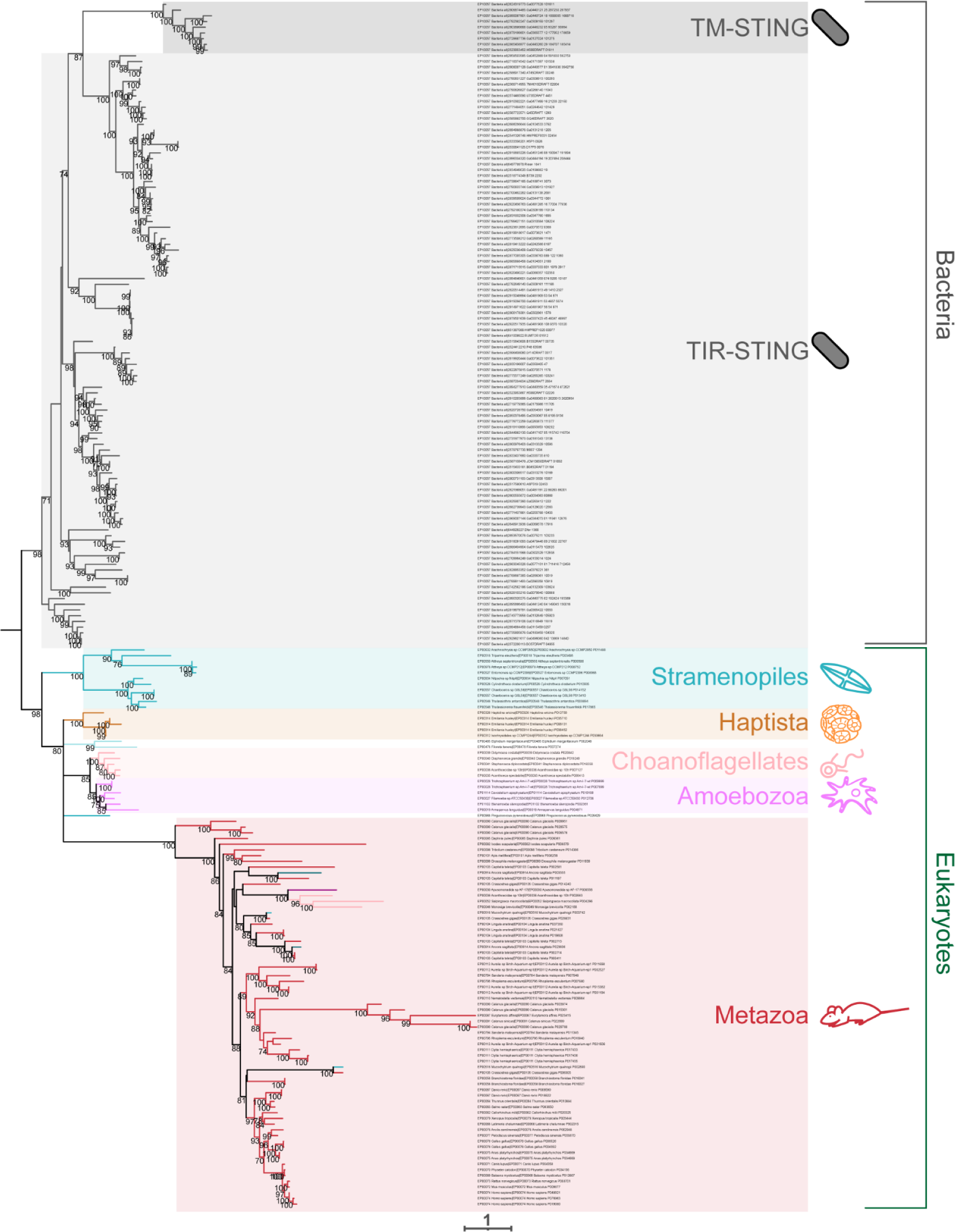
STING phylogenetic tree. Maximum likelihood phylogenetic tree of hits from iterative HMM searches for diverse eukaryotic STING domains. Scale bar represents the number of amino acid substitutions per position in the underlying MUSCLE alignment. Ultrafast bootstrap values calculated by IQtree at all nodes with support >70 are shown. Branches with support values <70 were collapsed to polytomies.

**Supplementary Figure 6:**
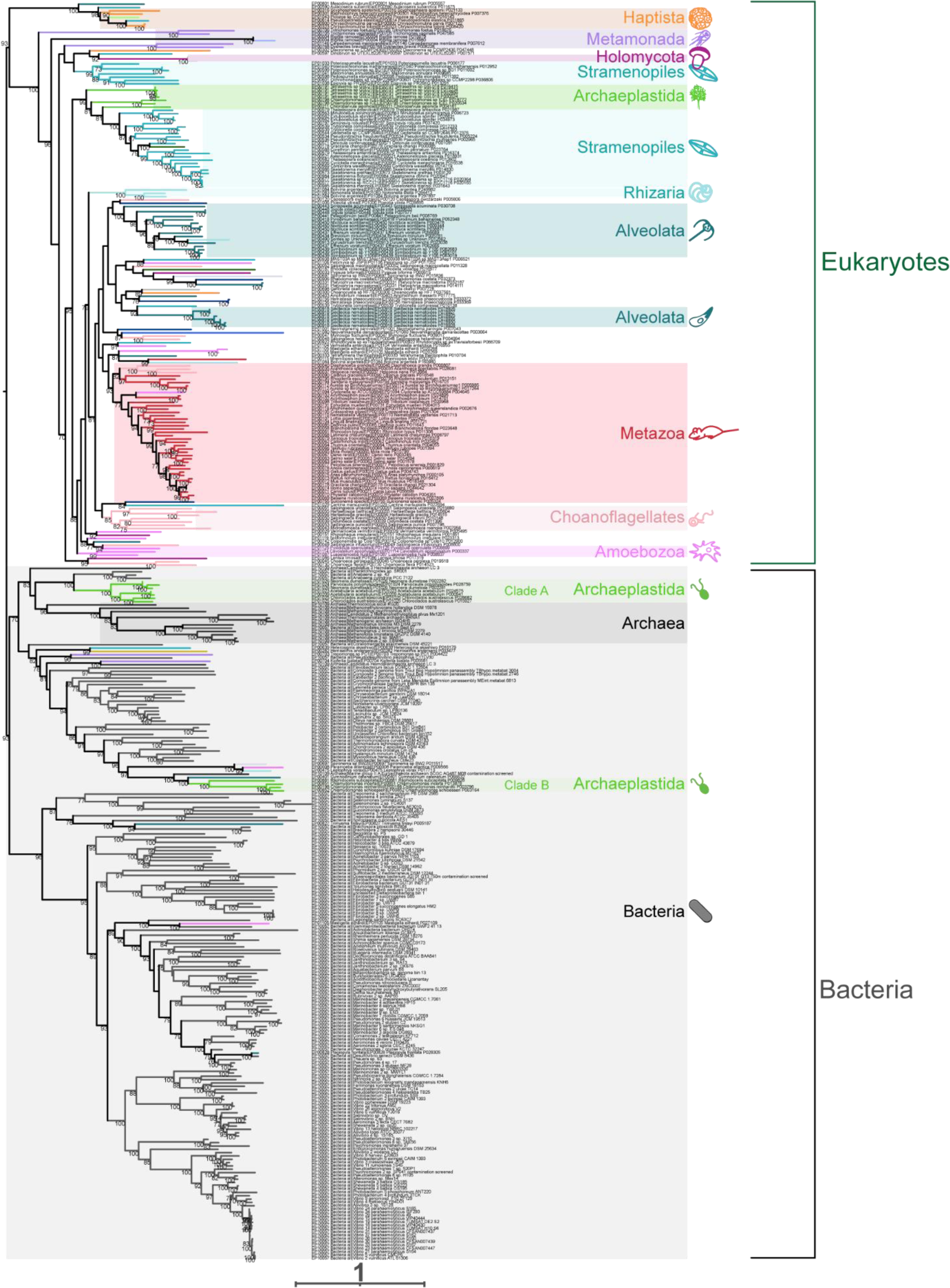
Viperin phylogenetic tree. Maximum likelihood phylogenetic tree generated by IQtree of hits from iterative HMM searches for diverse eukaryotic Viperins. Scale bar represents the number of amino acid substitutions per position in the underlying MUSCLE alignment. Ultrafast bootstrap values calculated by IQtree at all nodes with support >70 are shown. Branches with support values <70 were collapsed to polytomies.

**Supplementary Figure 7:**
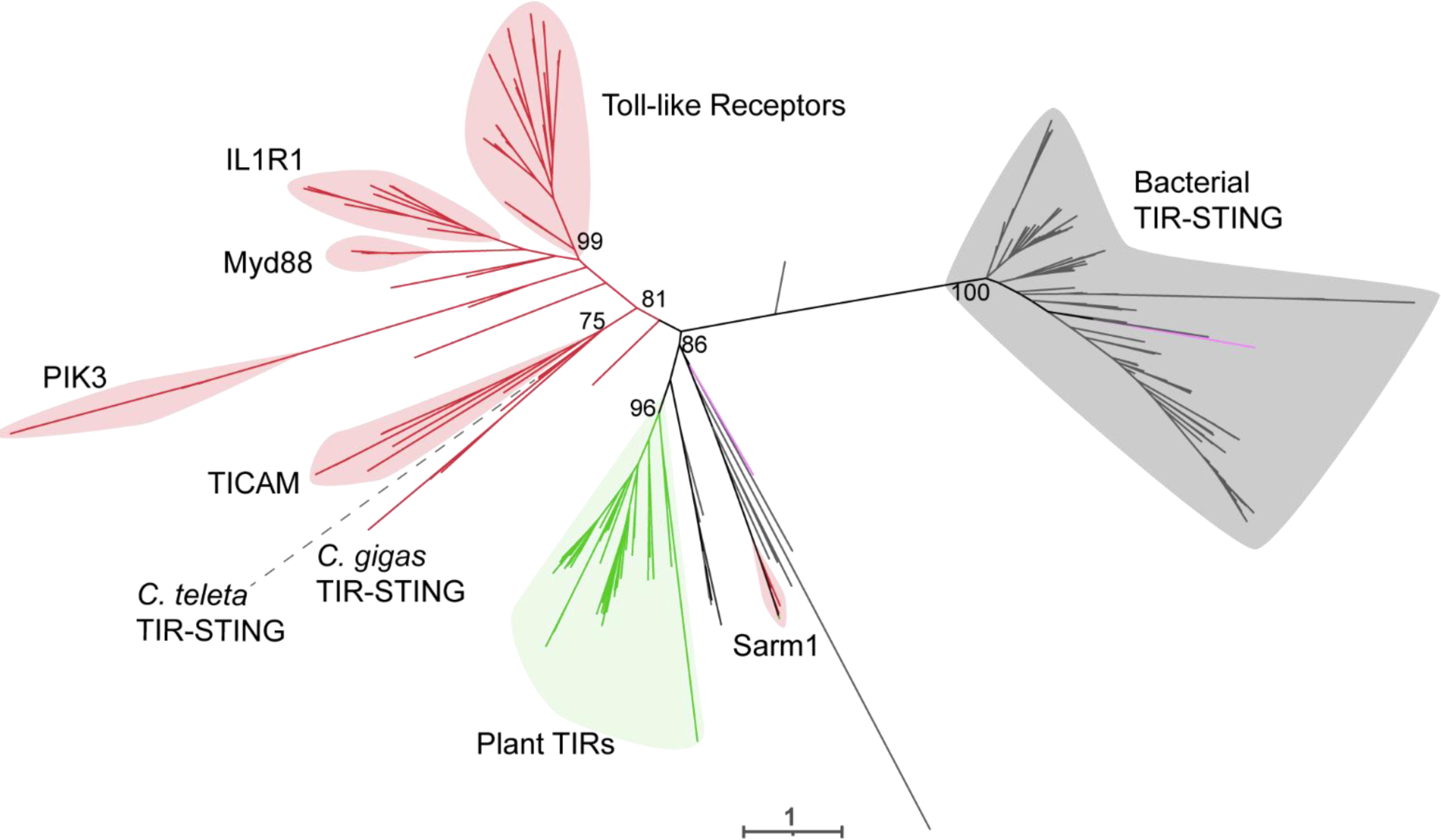
TIR domain of *Crassostrea gigas’* TIR-STING is closely related to metazoan TIR domains. Unrooted maximum likelihood tree of diverse TIR domains. Scale bars on the phylogenetic tree represent the number of amino acid substitutions per position in the underlying MUSCLE alignment. Ultrafast bootstrap values calculated by IQtree at key nodes are shown.

## Supplementary Data

**Supp. Data Fasta 1: CD-NTase** Fasta file with all CD-NTase amino acid sequences analyzed.

**Supp. Data Fasta 2: STING** Fasta file with all STING amino acid sequences analyzed.

**Supp. Data Fasta 3: Viperin** Fasta file with all Viperin amino acid sequences analyzed.

**Supp. Data Table 1: Hmmscan excel file** Hmmscan data for each CD-NTase, STING, and Viperin protein sequence.

**Supp. Data Tree 1: CD-NTase** Newick file of maximum likelihood phylogenetic tree generated from a MUSCLE alignment with IQtree. Newick file is used in Fig. 2 and Supp. Fig. 3 and 4. Node support values calculated from ultrafast bootstraps.

**Supp. Data Tree 2: STING** Newick file of maximum likelihood phylogenetic tree generated from a MUSCLE alignment with IQtree. Newick file is used in Fig. 3 and Supp. Fig. 3 and 5. Node support values calculated from ultrafast bootstraps.

**Supp. Data Tree 3: Viperin** Newick file of maximum likelihood phylogenetic tree generated from a MUSCLE alignment with IQtree. Newick file is used in Fig. 4 and Supp. Fig. 3 and 6. Node support values calculated from ultrafast bootstraps.

